# An autochthonous model of lung cancer in the Naked Mole-Rat (*Heterocephalus glaber*)

**DOI:** 10.1101/2023.08.28.555115

**Authors:** Alyssa Shepard, Scott Troutman, Sany Hoxha, Daniel Lester, Walid Khaled, Ewan St. John Smith, Thomas Park, Rochelle Buffenstein, Dongliang Du, Mingxiang Teng, Christine Crish, Kenneth Y. Tsai, Elsa R. Flores, Andrea Ventura, Joseph L. Kissil

**Affiliations:** Department of Molecular Oncology, The H. Lee Moffitt Cancer Center and Research Institute, FL, USA; Department of Pharmacology, University of Cambridge, Cambridge, UK; Department of Biological Sciences, University of Illinois Chicago, IL, USA; Department of Bioinformatics, The H. Lee Moffitt Cancer Center and Research Institute, FL, USA; Department of Pharmaceutical Sciences, Northeast Ohio Medical University, OH, USA; Departments of Anatomic Pathology and Tumor Biology, The H. Lee Moffitt Cancer Center and Research Institute, FL, USA; Cancer Biology and Genetics Program, Memorial Sloan Kettering Cancer Center, New York, NY, USA

**Keywords:** Naked Mole-Rat, Cancer, tumor initiation, CRISPR

## Abstract

Studies on cancer resistance in the naked mole-rat (NMR) have generally failed to interrogate possible resistance mechanisms in a physiological context. Here, we provide evidence that the NMR presents as a novel model of tumor initiation. We developed an endogenous lung cancer model in NMRs, driven by an oncogenic Eml4-Alk fusion protein introduced through CRISPR-mediated genome editing. While this is sufficient to drive tumorigenesis in mice, the development of progressive disease in NMRs required the additional loss of key tumor suppressors. Our results show that tumor initiation in NMRs more closely recapitulates that of human tumors. This suggests that the proposed “resistance” of NMRs to cancer development may stem from tumor initiation events that are likely to be comparable to the mechanisms in human cells.

**One-Sentence Summary:** Tumor development in the cancer-resistant naked mole-rat more accurately represents the tumor initiation process in humans.

## Introduction

Naked mole-rats (NMRs, *Heterocephalus glaber*) have emerged as a popular model for the study of cancer resistance and aging (*1, 2*). This spike in popularity can be attributed, in part, to their incredible lifespan and seldom reported cases of malignant disease (*3–7*). Over the past decade, NMRs have further cemented their reputation as a cancer-resistant species through a plethora of studies proposing multiple biological mechanisms that could contribute to their longevity and healthy lifespan (*8*). These mechanisms are varied, including early cell contact inhibition (*9*), enhanced DNA repair mechanisms and genome stability (*10–12*), high tolerance to oxidative stress (*13, 14*), and reduced inflammatory responses (*15*). Despite the steady increase in NMR-focused studies, many of these proposed mechanisms are limited to *in vitro* analyses and have yet to be truly tested in a physiological context through rigorous *in vivo* studies. Additionally, to date, there are no genetically engineered models of disease established in NMRs. To address these shortcomings, we sought to develop a first of its kind endogenous cancer model in the NMR.

Evidence of spontaneous tumors in NMRs has previously been reported (*5–7*), therefore we hypothesized that NMRs would develop tumors, given the appropriate initiating events, albeit at a delayed rate when compared to mice. The specific biological processes that may influence this delayed tumorigenesis are yet to be determined, but we hypothesized that NMRs may have different requirements for cellular transformation. It is well established that there are central differences in cellular transformation of rodent cells compared to human cells. In mice, the overexpression of one or two cooperating oncogenes is enough to lead to aberrant cell growth and subsequent transformation. In contrast, oncogene activation alone is not sufficient in human cells (*16–19*), and additional events are required, including the loss of cell cycle and cell death regulators, such as the tumor suppressors p53 and pRb, and the expression of telomerase (*16–19*).

To test our hypothesis, we developed a model of lung cancer using CRISPR-Cas9 to induce an oncogenic inversion, which would result in a fusion protein of *Eml4* and *Alk*. This chromosomal inversion is present in 4-6% of human lung adenocarcinoma (LuAD) cases and is a strong oncogenic driver in mice (*20–22*). In a previously established *Eml4-Alk* mouse model, the induction of the translocation by CRISPR-Cas9 was sufficient to drive significant tumor development with 100% penetrance within three months of infection (*21*). We utilized this approach to generate the *Eml4-Alk* inversion in NMRs and assess tumorigenesis in the lung.

### Validation of the CRISPR-Cas9 model and adenoviral infection in naked mole-rats

Given that an *in vivo* NMR model has not been developed to-date, the assessment of CRISPR- Cas9 functionality and suitability of an adenoviral-based delivery approach is critical. The *Eml4* and *Alk* alleles are located on the same chromosome in both mice and NMRs (Fig. S1a). We therefore adapted the dual-guide CRISPR-Cas9 strategy used in the murine model for editing of the NMR genome (Fig. S1b).

To first assess the suitability of an adenoviral-based delivery system, we infected NMR lungs with an adenovirus containing GFP (Ad-GFP) via intra-nasal instillation, collected the lungs 72 hours post infection, and analyzed the dissociated cells via flow cytometry (Fig. S2a).

Additionally, we tested a dual-adenoviral infection using Ad-GFP and Ad-RFP, to ensure the dual-infection leads to a sufficient subset of double positive NMR lung cells (Fig. S2b). Following confirmation of successful infection, we tested the CRISPR-Cas9 adenovirus in NMR cells. We infected immortalized NMR skin fibroblasts (*23*) (ISF) *in vitro* or NMR lungs *in vivo* with the *Eml4-Alk* CRISPR virus (Ad-EA) and collected the lungs 72 hours post infection. Genomic DNA was extracted and recombination assessed by PCR. This analysis confirmed that these infections led to the successful generation of the *Eml4-Alk* inversion in ISFs and NMR lungs (Fig. S2c).

### Functional Eml4-Alk fusion protein leads to a growth advantage in NMR skin fibroblasts

After confirming that the designed genomic inversion event occurred, we next confirmed whether there was a functional Eml4-Alk fusion protein in our infected cells by using a cell proliferation assay. Towards this goal we established a set of NMR skin fibroblast cell lines, both primary (PSF) and immortalized (ISF), that express the *Eml4-Alk* mRNA fusion, as confirmed by RT-PCR (Fig. 1a). The growth of the *Eml4-Alk* cell lines (*Eml4-Alk***^+^**) was then compared to control cells. The presence of the *Eml4-Alk* inversion resulted in a significant growth advantage for the *Eml4-Alk***^+^** over control cells, as shown in both PSF and ISF (Fig. 1b and Fig. S3). Treatment with the FDA-approved Alk inhibitor crizotinib impaired this growth advantage in the *Eml4-Alk***^+^**immortalized cell lines, suggesting that these cells do have a functional, constitutively active Eml4-Alk protein that contributes to a growth advantage in these cells (Fig. 1c).

**Fig. 1.**
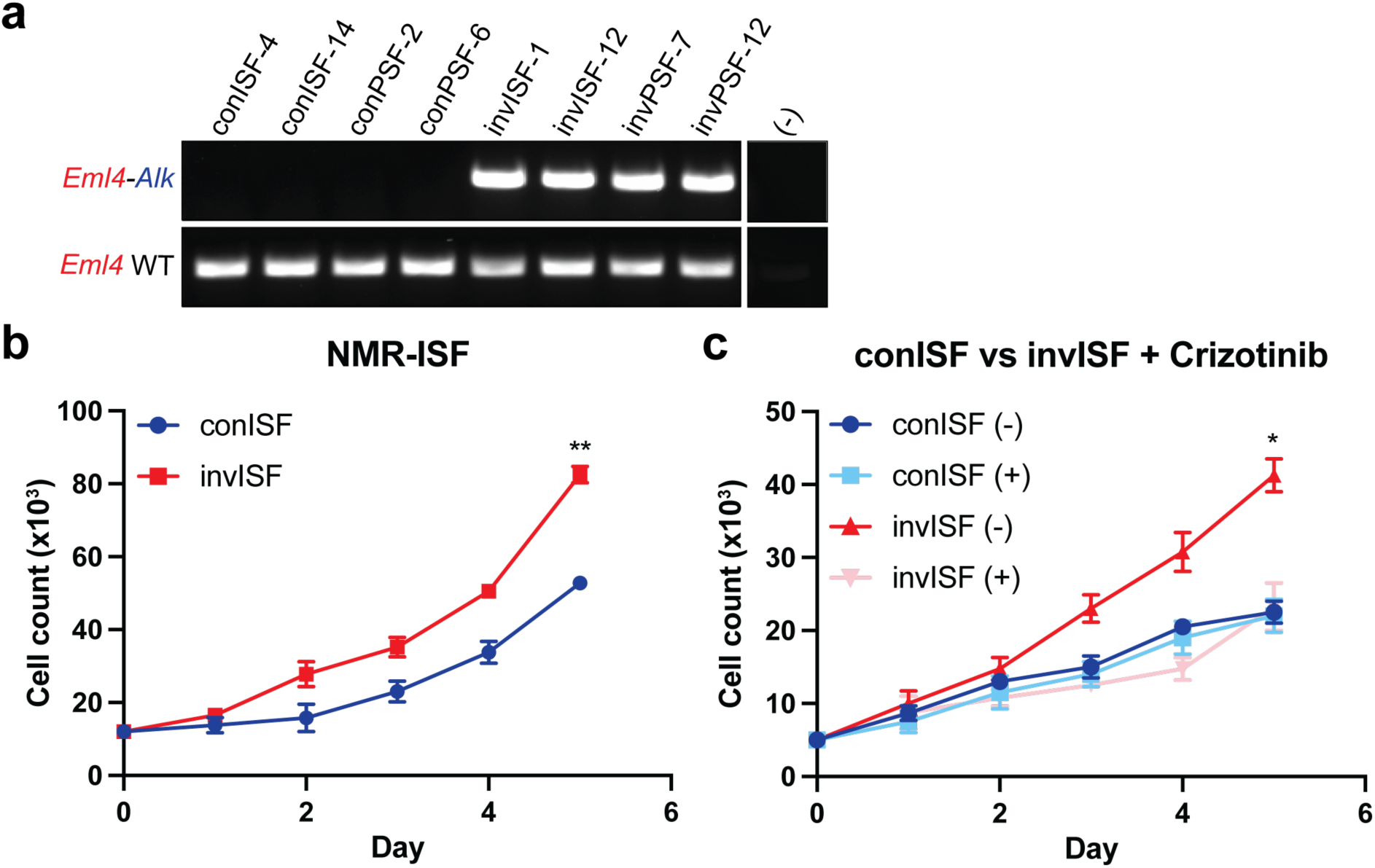
Effect of *Eml4-Alk* expression in NMR skin fibroblasts. (**a**) RT-PCR detection of the *Eml4-Alk* inversion. RNA was extracted from NMR skin fibroblast cell lines, both immortalized (ISF) and primary (PSF). ‘Con’ indicates control cell lines and ‘inv’ indicates cell lines with *Eml4-Alk*. Two independent cell lines were generated for each through clonal expansion. (-) indicates no template negative control. (**b**) Growth assay for NMR-ISF cell line. Cells were plated in triplicate in a 12-well plate, 10,000 cells per well and counted over five days. Growth assays shown are representative of three independent experiments. Cell lines chosen are representative of one of two cell lines made with the *Eml4-Alk* inversion. Two-way ANOVA determined the interaction of time and genotype to have a statistically significant effect on cell count (F(5, 20) = 30.03; p-value < 0.0001). Genotype alone also had a statistically significant effect on cell count (p-value < 0.0001). Multiple comparisons determined cell counts on day 5 to be significantly different (**, p-value = 0.0016). (**c**) Growth assay was prepared as described in (**b**), starting with 5,000 cells per well. Cells were treated with 0.25μM Crizotinib. Three-way ANOVA determined the interaction of time, genotype, and treatment to have a statistically significant effect on cell count (F(5, 40) = 12.85; p-value < 0.0001). The interaction of genotype and treatment also had a statistically significant effect on cell count (F(1, 8) = 406.7; p-value < 0.0001). Multiple comparisons determined cell counts for invISF (-) and invISF (+) on day 5 to be significantly different (*, p-value = 0.0397).

### Induction of the *Eml4-Alk* inversion does not induce tumorigenesis in NMR lungs

NMRs were next infected intra-nasally with Ad-EA or a control CRISPR virus (Ad-con). Lungs from both infection groups were collected at three months post-infection and preserved via FFPE processing. The three-month timepoint was chosen based on previous results in the mouse model in which all animals infected with an inversion-inducing adenoviral vector displayed full blown lung adenocarcinoma (*21*). Upon examination, none of the control NMRs showed evidence of any tumor development as visualized by H&E (Fig. 2a-b). H&E staining showed normal pathology consistent with healthy lungs. The *Eml4-Alk* infections recapitulated what we observed in the control infections. The NMRs displayed no evidence of pre-cancerous pathology at a 3-month timepoint (Fig. 2c-d). An additional group of NMR infections were carried out to a 6-month timepoint, again with no evidence of tumor development (Fig. S4a-b).

**Fig. 2.**
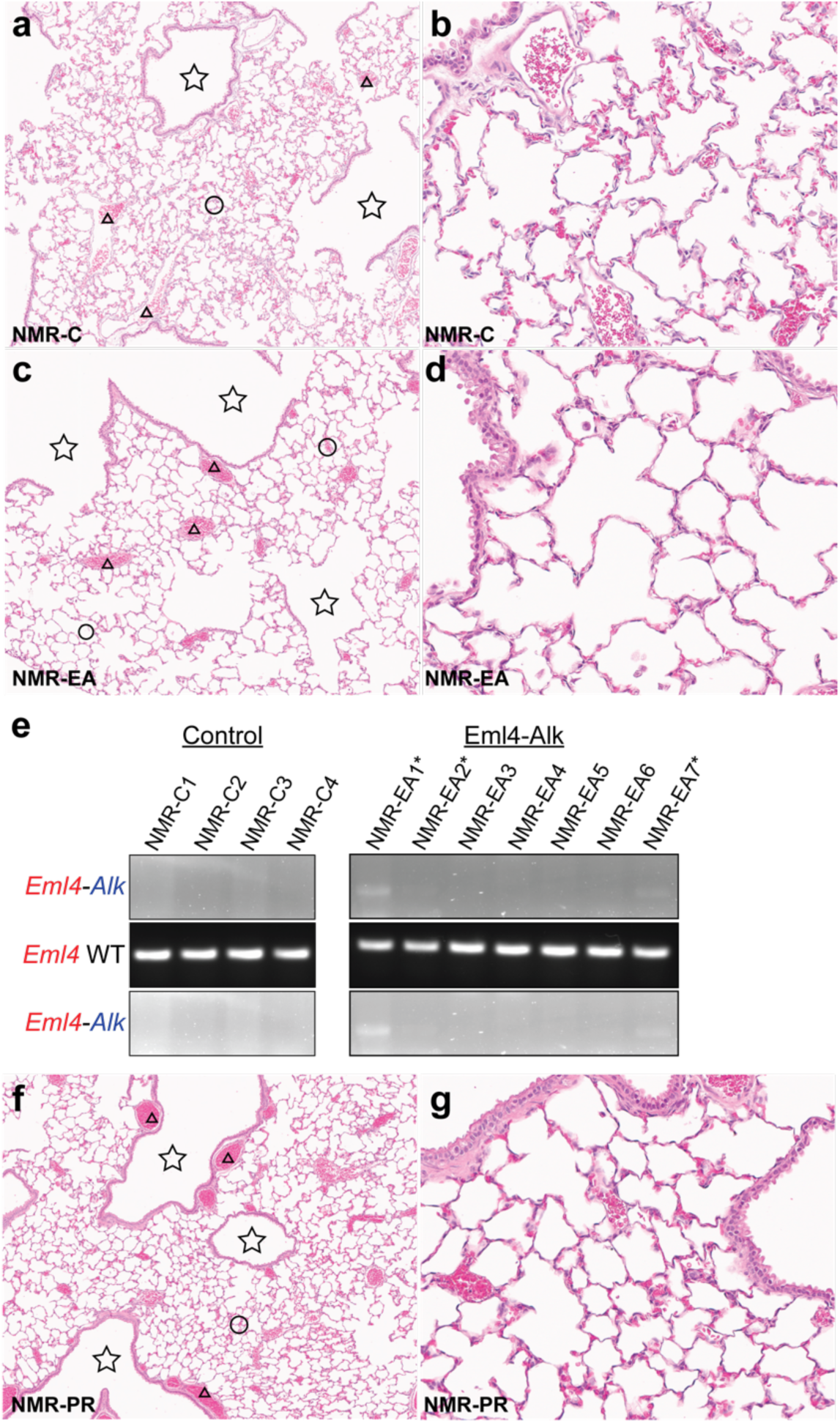
Control and *Eml4-Alk* infections at a 3-month timepoint. Infected NMR lungs were collected, preserved through FFPE, and processed for H&E staining. Representative images of lungs from NMR control (NMR-C) groups (**a-b**; n=4), *Eml4-Alk* (NMR-EA) groups (**c-d**; n=7), and *Tp53/Rb1* groups (**f-g**; n=5) at a 3-month timepoint. Star = airway; circle = normal lung tissue; triangle = blood vessel. (**a**) Representative image for control infection, 50X magnification. (**b**) Representative image for control infection, 200X magnification. (**c**) Representative image for *Eml4-Alk* infection, 50X magnification. (**d**) Representative image for *Eml4-Alk* infection, 200X magnification. (**e**) Genomic PCR analysis of DNA extracted from FFPE tissues of control and *Eml4-Alk* infections. *Eml4-Alk* gels are shown overexposed in bottom row. Asterisk indicates detectable *Eml4-Alk*. (**f**) Representative image for *Tp53/Rb1* infection (NMR-PR), 50X magnification. (**g**) Representative image for *Tp53/Rb1* infection (NMR-PR), 200X magnification.

We extracted DNA from FFPE sections from the three month infections and analyzed them via genomic PCR (Fig. 2e). From these, none of the control infections had detectable levels of the *Eml4-Alk* inversion, while a subset of NMR lungs from the Ad-EA infections did present with detectable levels of *Eml4-Alk* (3/7). These data suggest that the presence of the *Eml4-Alk* inversion alone is not sufficient to induce tumorigenesis in NMR lungs.

### Simultaneous loss of *Tp53* and *Rb1* does not induce tumorigenesis in NMR lungs

Unlike in mice, where the expression of a strong oncogenic driver can lead to widespread tumor growth in the lung, the induction of the *Eml4-Alk* inversion did not lead to tumorigenesis in NMRs. This is in line with our initial hypothesis, that NMRs have requirements for cellular transformation that differ from mice. To further investigate, we added two additional driving events: the loss of the tumor suppressors p53 and pRb. *Tp53* and *Rb1* mutations are common co-alterations in ALK^+^ NSCLC patients (*24, 25*). Additionally, *Tp53* and *Rb1* mutations are commonly used to drive tumorigenesis in small cell lung cancer (SCLC) models in mice (*26–28*).

We next developed an additional CRISPR-Cas9 adenovirus to target both *Tp53* and *Rb1* (Ad-PR). This adenovirus was modeled after the same dual-guide CRISPR system we used for *Eml4-Alk*, with one guide RNA (sgRNA) targeting *Tp53* and one for *Rb1* to create inactivating mutations and/or indels in coding exons. The generation of these mutations was confirmed by infecting NMR PSF with Ad-PR, collecting DNA after 72 hours, PCR amplifying the target region, and sequencing (Fig. S5). Following this confirmation, NMRs were infected with Ad-PR through intranasal instillation. Lungs infected with Ad-PR showed no signs of tumor development at a 3- month timepoint, confirmed by H&E staining (Fig. 2f-g).

In addition to the first-generation Ad-PR, we made a set of second-generation dual guide adenoviruses for *Tp53* (Ad-p53-d) and *Rb1* (Ad-Rb1-d), which result in large genomic deletions of the respective genes. The purpose of this was to make identification of *Tp53* and *Rb1* mutations simpler, by following the same PCR-based identification system we have used for the *Eml4-Alk* inversion. We assessed the virus *in vivo* at 72-hour post-infection to confirm that these deletions are detectable in NMR lungs (Fig. S6a). Previously, we validated that NMR lungs would have a sufficient percentage of cells that are double-positive upon infection with two adenoviruses (Fig.S2b). These second-generation CRISPR viruses were then used in additional dual-infections with the Ad-EA virus to determine if combinations of *Eml4-Alk/Tp53* or *Eml4-Alk/Rb1* mutations would lead to tumor formation. As was the case with the *Eml4-Alk* only infections, the NMR lungs appear normal and tumor-free at a 15-week timepoint (Fig. S6b-e).

### The combined expression of *Eml4-Alk* and loss of both *Tp53* and *Rb1* is required to lead to significant lung tumor burden in NMRs

Upon confirming that combinations of *Eml4-Alk* expression and mutations in either *Tp53* or *Rb1* did not lead to tumorigenesis, we assessed whether the combined effect of all three driving events could drive tumor development. To do so, we co-infected NMRs (n=10) with the Ad-EA and Ad- PR viruses. From these infections, a subset of our NMRs (3 of 10) developed significant tumor growth at around a 15-week timepoint. The morphology of the tumors was consistent with pleomorphic carcinoma, mostly comprised of adenocarcinoma with areas of spindle cell and giant cell carcinoma (Fig. 3a-e). IHC staining was largely heterogenous, with distinct areas of positive staining for the markers SPC-C, CK14, and NKX2-1 (Fig. 3f-h). Most tumors stained strongly with Ki67, meaning the tumors were highly proliferative, with an estimated Ki67 index of 80% (Fig. 3i). Overall, our IHC analysis was sufficient to provide a diagnosis of pleomorphic carcinoma. As is seen in human pleomorphic carcinoma, our tumors are likely of epithelial origin, but have areas of dedifferentiation into an epithelial-mesenchymal transition.

**Fig. 3.**
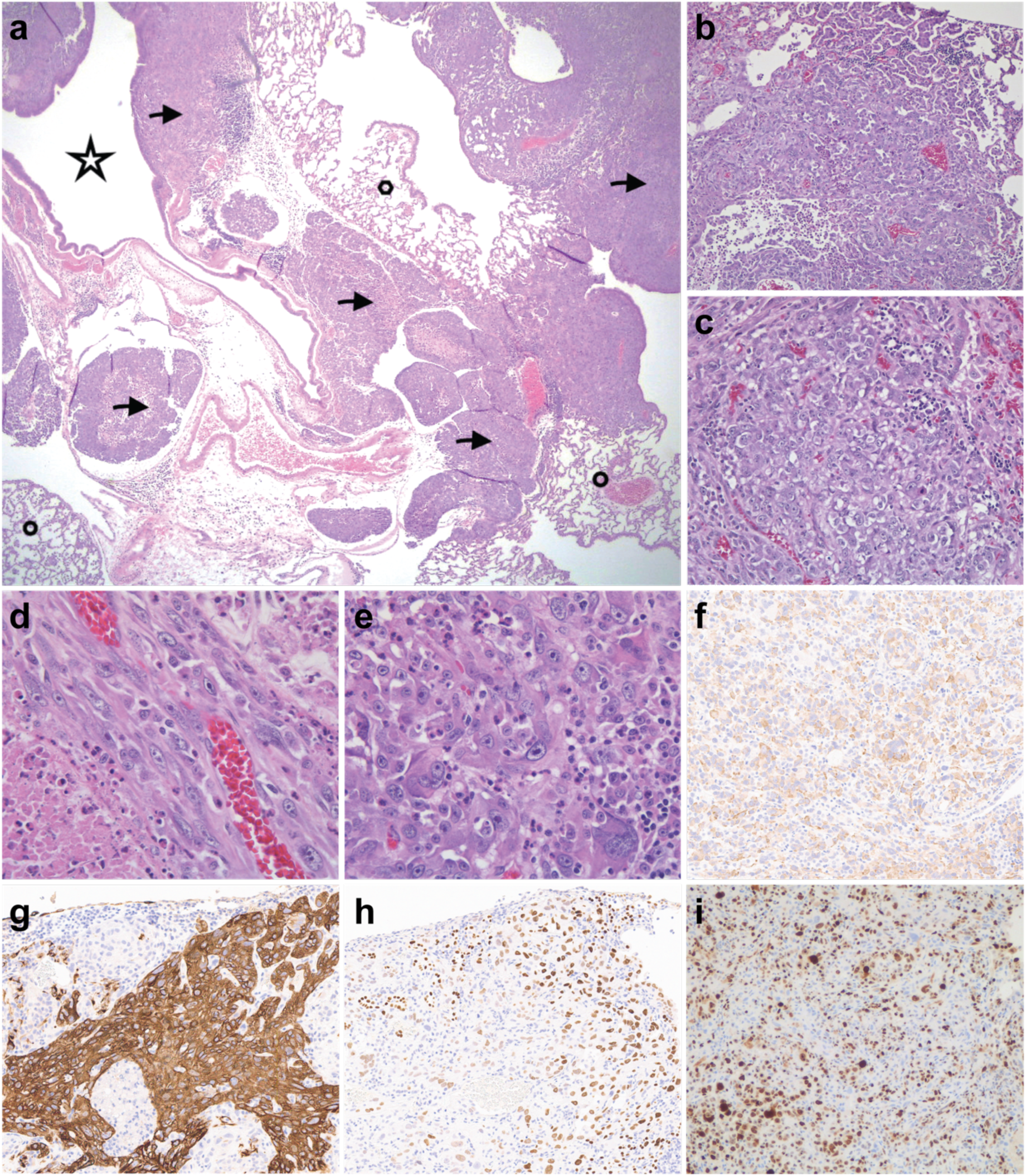
Histology and IHC of NMR lungs from dual-infection group with significant tumor burden. NMR lungs from infections were collected, preserved through FFPE, and processed for H&E and IHC staining. Representative images of lungs from the NMR dual infection group (Ad-EA/Ad-PR). (**a-e**) H&E staining. (**a**) Overview of NMR lung. Star = airway; circle = normal lung tissue; arrows = adenocarcinoma nodules. 25X magnification. (**b-c**) Solid adenocarcinoma nodules. Tumor cells are large and round with pleomorphic nuclei, prominent nucleoli, and large cytoplasms. (**b**) 200X magnification. (**c**) 400X magnification. (**d**) Pleomorphic spindle/fusiform cells in a carcinomatous embolism. 400X magnification. (**e**) Giant tumor cells with polymorphic and multi-nucleated nuclei with immune cell infiltration. 400X magnification. (**f**) SPC-C. 200X magnification. (**g**) CK14. 200X magnification. (**h**) NKX2-1. 200X magnification. (**i**) Ki67. 200X magnification.

Mutational analysis of these tumors was conducted by isolating individual tumor regions using laser capture microdissection (LCM) of FFPE tissues (Fig. 4a). DNA was extracted from these tumors and used for genomic PCR to detect the *Eml4-Alk* inversion or targeted deep sequencing of *Tp53* and *Rb1* to detect mutations or small deletions (indels). This analysis revealed that all the tumors from which DNA was successfully isolated contain the *Eml4-Alk* inversion (Fig. 4b). Furthermore, deep sequencing analysis confirmed that all isolated tumors also contained mutations in *Tp53* and *Rb1* (Fig. 4c). The most common *Tp53* and *Rb1* mutations, which were identified in 100% of the tumors, were disruptive frameshift mutations which have a high probability of leading to an inactive protein. Most tumors contained additional mutations in these two genes which could further disrupt function. The mutational profiles of these tumors, along with the results of the previous infection experiments we conducted, suggests that all three driving events are necessary to drive formation of tumors in NMR lungs.

**Fig. 4.**
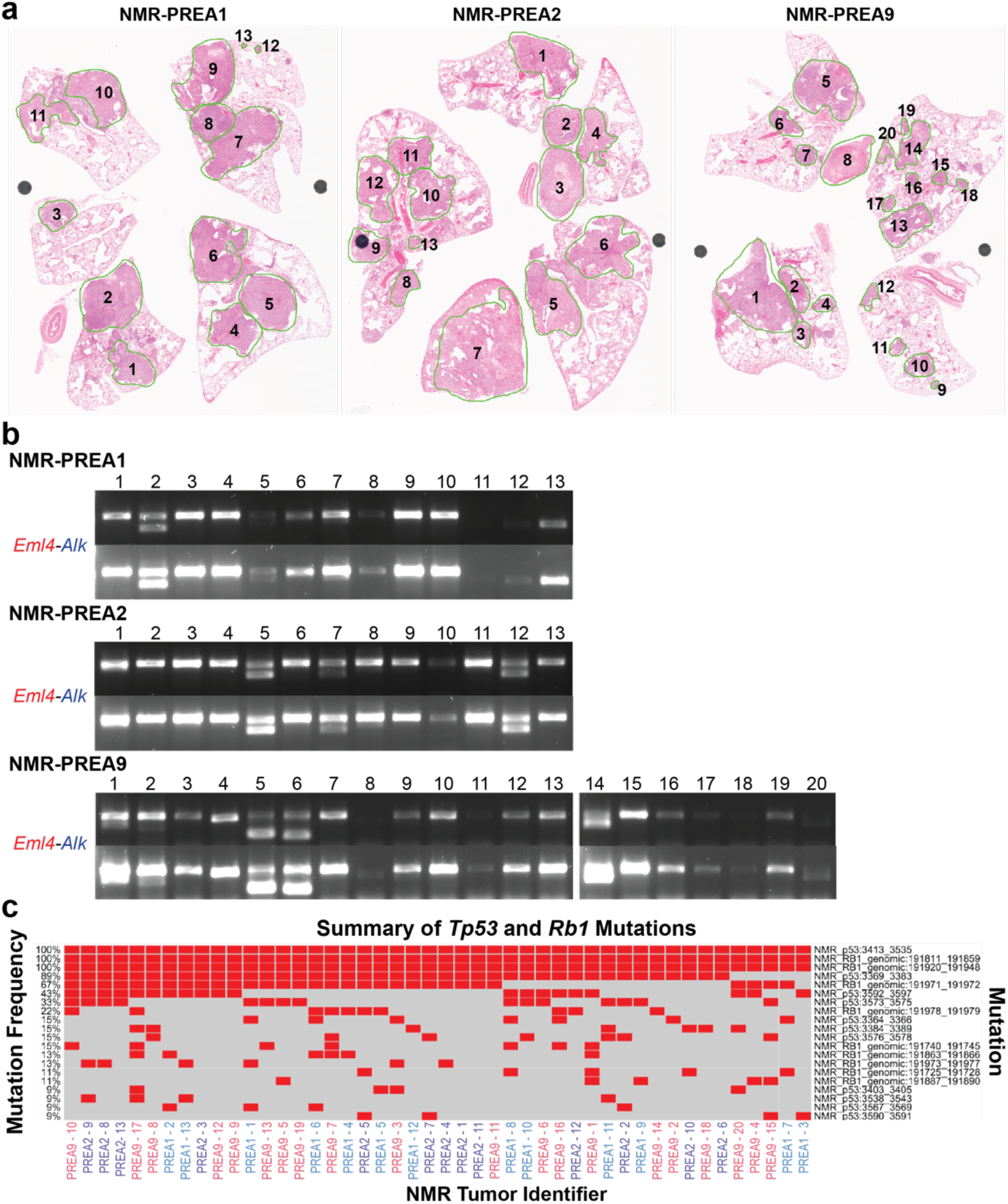
LCM strategy and subsequent mutation analysis. (**a**) Whole-slide images of the NMR lungs with significant tumor burden. Tumors were isolated as shown by the green lines. Number indicates tumor identification. Images were scanned at 20X magnification. (**b**) DNA was extracted from LCM-captured tumors. Genomic PCR analysis shows the presence of the *Eml4-Alk* inversion. Tumor identification numbers are indicated above the gels. Low and high exposures shown. (**c**) Top 20 mutations identified in *Tp53* and *Rb1* with an allelic frequency of 0.01 or greater. The percent frequency of each mutation is indicated on the left Y-axis. Individual tumors are shown on the X-axis, color coordinated by the NMR the tumor was taken from: NMR-PREA1, blue; NMR-PREA2, purple; NMR-PREA3, red. Red boxes indicate which mutations each tumor has. Mutation type is indicated on the far right.

## Discussion

As previous studies suggested that NMRs are resistant to the development of tumors, we set out to assess this *in vivo* using an approach previously used to generate a model of *Eml4-Alk* driven lung adenocarcinoma in mice. We utilized CRISPR-mediated genome editing to induce this inversion in somatic cells of the NMR lung. Our data indicate that while we could generate the designed inversion, which resulted in expression of an active *Eml4-Alk* fusion protein, this alone was insufficient to drive tumorigenesis in the NMR lungs. This is in contradiction to what is seen in mice, where *Eml4-Alk* alone leads to rapid, widespread tumor growth (*21*). Based on these findings, we hypothesized is that NMRs likely require additional “hits” to lead to cellular transformation *in vivo*, as suggested to be the case in human cells.

Human cells require more than the activation of an oncogene to lead to cellular transformation. Additional events include the loss of cell cycle regulators, cell death regulators, and the expression of telomerase (*16, 18*). To assess whether NMRs had similar transformation mechanisms, we targeted the tumor suppressor genes *Tp53* and *Rb1*. We first assessed these alone or in combination and found that, similar to the *Eml4-Alk* infections, infections with our Ad-PR to induce *Tp53* and *Rb1* mutations also did not lead to tumor development. Furthermore, the combinations of *Eml4-Alk/p53* or *Eml4-Alk/Rb1* mutations did not lead to tumorigenesis. Previous literature has suggested that NMR cells can be transformed in culture with the simultaneous loss of p53 and pRb (*9*). This has been contradicted in a subsequent study that observed cellular crisis and a cessation of growth with loss of p53 and pRb (*29*), which our *in vivo* data more closely aligns with. We have shown here that the loss of these critical tumor suppressors fails to lead to tumor development. Similarly, using the combination of the expression of an oncogene and loss of a single tumor suppressor we were unable to observe any lesions in the NMR lungs, suggesting the combinations of these events are insufficient in inducing tumorigenesis. Tumorigenesis in the NMR was only observed in NMR infected with the combination of the *Eml4-Alk* inversion and both *Tp53* and *Rb1* mutations. Overall, our model had a penetrance of around 30%, likely due to the ability of the dual infections to target the relevant tumor initiating cells simultaneously.

The tumors that developed in the NMR lungs were large and heterogenous and likely indicative of late-stage disease. Significantly, the tumors were identified as pleomorphic lung carcinoma, a rare subtype of NSCLC for which no mouse models are currently available. Given the rarity of this form of NSCLC, there are not many studies with published data sets compiled from a limited number of patients, and obtaining access to these has proved challenging. However, there is evidence that the majority of pleomorphic lung tumors have a mutation in *Tp53*, along with additional driver mutations such as *Kras*, *EGFR*, and *Alk* fusions (all identified as *Eml4-Alk* fusions), with *Alk* mutations being relatively rare (*30, 31*). The driving mutations are generally mutually exclusive, an observation consistent with other subtypes of NSCLC (*32*). Mutations in *Rb1* were also uncommon, but not mutually exclusive with the *Tp53* alterations (*30, 31*). Future comparative analysis using human data sets, when these become available, would be beneficial in identifying similarities between the NMR tumors and human disease.

With this diagnosis, we have not only provided evidence of critical requirements for tumorigenesis in NMRs, but also generated a genetic model that could fill a gap in the NSCLC field with further development. Importantly, we have shown that NMRs may more closely recapitulate human tumor initiation and may provide a better representation for tumor initiation. Continuing studies on NMRs and the generation of additional *in vivo* models with different driving events will be essential for confirming this hypothesis. Our observations do not necessarily discredit the claims that NMRs are resistant to cancer, rather, confirm a difference in the susceptibility to development of cancer between mice and NMRs. An important next step is distinguishing whether this susceptibility is impacted by variations in cell autonomous versus non-cell autonomous functions. Whether the NMR immune system, for example, plays a role in the unique tumor initiating events observed in the NMR remains to be determined, and will require the development of species-specific reagents.

## Acknowledgements

We thank Dr. Elisabeth Brambilla for providing the diagnosis of the lung tumors and consulting on the H&E and IHC images.

## Author contributions

Conceptualization: AS, ST, SH, AV, JLK

Methodology: AS, ST, SH, AV, ERF, KT

Investigation: AS, ST, SH, DL, ERF

Resources: WH, ESTJ, TP, RB, CC, AV, ERF

Formal Analysis: DD, MT Funding acquisition: JLK

Supervision: JLK

Writing – original draft: AS, JLK

Writing – review & editing: AS, JLK

## Data and materials availability

Deep sequencing data available upon request. All other data are available in the main text or in the supplemental materials.

