## Supplementary material for "An autochthonous model of lung cancer in the Naked Mole-Rat (*Heterocephalus glaber*)": Supp files

### Materials and Methods

**Cell culture.** NMR skin fibroblasts (primary and immortalized) were grown in Dulbecco's Modified Eagle's Medium (DMEM; Gibco) supplemented with 15% fetal bovine serum (FBS; Atlas Biologicals), 1% penicillin/streptomycin (Gibco), 1% non-essential amino acids (Gibco), 1% sodium pyruvate (Gibco), and 0.2% primocin (InvivoGen). Cells were maintained at 32°C, 5% $CO_2$ and 3% $O_2$. NMR primary fibroblasts were immortalized using HRasG12V and SV40LT (NMR-ISF cells). NMR skin fibroblasts were gifted by the Khaled and St. John Smith Labs (University of Cambridge, UK) (*23*).

**NMR colony maintenance.** NMRs were housed in inter-connected cages that mimic the underground tunnel systems in their natural habitat. Three to six cages were used per colony, depending on colony size, and connected with 2" polycarbonate pipes, secured by screws. Temperature was maintained at 82-90˚F and humidity maintained between 40-60%. General considerations from Yu et al. were taken for breeding, husbandry, and general colony management (*33*).

**Cloning.** Guide RNAs (sgRNAs) were cloned into a variety of CRISPR plasmids, including PX459 (Addgene #62988), PX333 (Addgene #64073), and pAd5 (Addgene #64072). Plasmids were digested in CutSmart Buffer (NEB) with either BbsI (NEB) or BsaI (NEB). sgRNAs were annealed using T4 Polynucleotide Kinase (PNK, NEB). Annealed sgRNAs were ligated into the digested plasmid using Ligation Mighty Mix (Takara). Ligation products were transformed into TOP10 competent E. coli (Invitrogen). Positive clones were maxi-prepped with the ZymoPure II Plasmid Maxiprep Kit (Zymo Research). All reactions were carried out according to the manufacturer's instructions.

**Adenovirus infections.** *In vitro* – NMR cells were plated 500,000 cells in a 10cm dish 24 hours prior to infections. Unless otherwise stated, 10μl of adenovirus was used for infections (1 x $10^6$

pfu – $1x10^8$ pfu depending on adenovirus titer). Infection media was left for 72 hours before being replaced by fresh media. Cells were then either collected for analysis or left to proliferate.

*In vivo* – Infection mix was prepared using MEM, 1mM $CaCl_2$, and $3 \times 10^{10}$ pfu total of the selected adenovirus(es) to a total volume of 50µl. Infected NMRs were non-breeders between ages 1 to 3. NMRs were placed under anesthesia (isoflurane) and infected via intranasal instillation in two doses, 25µl each. Following infections, NMRs were observed until recovery and placed back in their appropriate colonies. Infections with a control virus (Ad-con) confirmed there were no animals that displayed an adverse response or developed detectable pathological manifestations as a consequence of adenovirus exposure.

**Genomic PCR.** Genomic DNA was extracted using the DNA Mini Kit (Qiagen). FFPE DNA was extracted using the GeneRead FFPE DNA Kit (Qiagen). All PCRs were done using AmpliTaq Gold 360 PCR Master Mix (Inivtrogen). PCRs were run as 50µl reactions according to manufacturer's protocol. PCR products were run on 1% agarose gels at 100V for 30' – 40' or sent for Sanger sequencing (Eton Bioscience and Genewiz).

**RT-PCR.** RNA was extracted using the RNeasy Mini Kit (Qiagen). cDNA was transcribed from total RNA using the iScript cDNA Synthesis Kit (Bio-Rad). 1µg RNA was used for each reaction, unless starting material was scarce, where the maximum amount of RNA was run based on available material. PCRs were run as previously described, using 1µl of cDNA from the RT.

**Growth assay.** Cells were seeded in 24-well culture plates at 10,000 cells per well (unless otherwise specified), in triplicate, for a 5-day assay (15 wells total). Cells were manually counted with a hemocytometer around the same time every day. Media was changed every other day. Significance was calculated using mixed effects analysis or two- or three-way ANOVA, depending on the number of experimental and treatment groups (indicated in figure legends). Multiple comparison analysis was also conducted to compare individual experimental conditions. Calculations were conducted using Prism version 7.

**Histology**. Lungs were harvested at the specified time points, inflated by intratracheal injection with 10% formalin (for fixation) or PBS (for immediate analysis), incubated for 24h in 10% formalin, and then transferred to 70% ethanol for another 24h. Lungs were then embedded in paraffin for future analysis. For staining, tissue slices were cut from FFPE blocks and mounted on microscope slides. Lungs from all NMRs were stained with hematoxylin and eosin (H&E). Lungs were also processed for a variety of markers with immunohistochemistry (IHC). Antibodies and dilutions used for this project are listed in Table 2.3. All slides were scanned at 20X magnification using the Leica Aperio AT2. 2.3.3. We observed that NMR lungs may be more fragile than mouse lungs when it comes to inflation. Our H&E stains often show ruptured airways and alveolar tissue, along with ruptured blood vessels that lead to blood cells ubiquitously observed amongst the alveolar sacs. This is merely due to the collection process and not a consequence of the adenovirus infections.

**Laser capture microscopy.** Laser capture microscopy (LCM) was completed using the Arcturus XT Laser Capture Microdissection System. Individual tumors were identified from the scanned H&E slides. An unstained, 10μm thick section of FFPE tissue was mounted on PEN membrane slides (2μm; Leica). Once tissue was cut, forceps were used to pick out dissected tissue from the slide and place in 1.5mL microcentrifuge tubes for DNA or RNA extraction.

**Targeted deep sequencing.** *Library preparations* – Following PCR amplification, amplicons were sequenced using the Illumina Nextera XT kit following the manufacturer's protocol (Illumina). Briefly, following the quality control screening of 100ng of each amplicon on a 1% agarose gel, both amplicons for each sample were pooled equimolar and 1ng of input DNA was used for the Nextera tagmentation reaction, followed by PCR amplification and sample indexing. The libraries were purified, quantitated with the Qubit Fluorometer (ThermoFisher Scientific), and screened on the Agilent TapeStation 4200 using the TapeStation High-Sensitivity D1000 Screen Tape (Agilent Technologies). The final libraries were normalized, denatured, and sequenced on the Illumina MiSeq sequencer using a Nano flow cell to generate approximately 20,000 150-base read pairs per sample.

*Data analysis* – Raw sequencing reads were first trimmed with cutadapt 1.8.1 and aligned to the target genes with bwa 0.7.10-r789. Duplicate reads were removed with GATK 4.1.0.0. Variants were then called using Mutect2 based on tumor-only mode. Variant allele frequency no less than 0.01 was applied to selected confident SNPs and 98 InDels. SNPs and InDels that were found at consecutive loci within each sample or across samples were further merged to define highly mutated spots. The oncoplots were generated based on the appearance of the highly mutated spots using ComplexHeatmap. The selected variants (MuTect2 AF >=0.01) were annotated using tool SnpEff (Version 4.3t) based on genome information (GCF_000247695.1_HetGla_female_1.0_genomic.gff).

**Flow cytometry.** Lungs were collected from NMRs and dissociated into single cells by digestion at 37˚C for 45 minutes with agitation in digestion buffer (Advanced DMEM/F12 (Gibco), 50 μg/mL penicillin/streptomycin (Gibco), 1 mg/mL collagenase (Sigma), 40 U/mL DNase I (Roche), 5μM HEPES (Gibco), 0.36mM $CaCl_2$). Cell suspension was washed twice in PBS and filtered through 0.7μm filters to remove large chunks and promote single cell suspension. Cells were incubated in red cell lysis buffer (ACK Buffer; Thermo) for 5 minutes at room temperature. Cells were washed twice more in PBS and resuspended in flow buffer (PBS, 0.1% BSA, 2mM EDTA). Cells were analyzed using a BD LSRII flow cytometer.

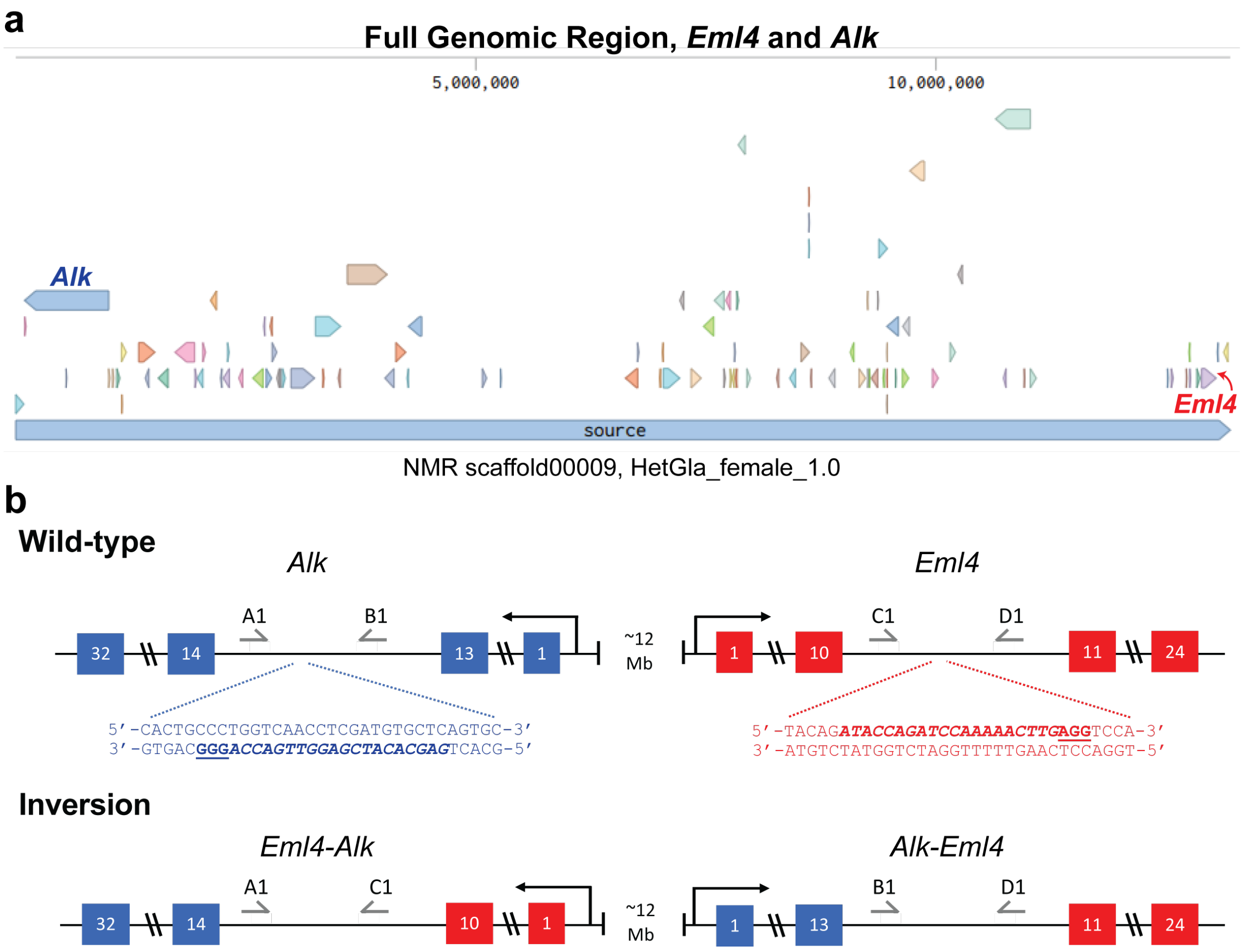


**Fig. S1. Genomic context of *Eml4* and *Alk* in the NMR and strategy for the *Eml4-Alk* inversion. (a)** Schematic of the unplaced NMR scaffold00009 from the NMR whole genome sequence (HetGla_female_1.0). Genomic sequences of *Alk* and *Eml4* are labeled. Other arrows indicate additional genes along the scaffold. (**b**) Schematic representation of the *Eml4-Alk* inversion. Exon numbers are indicated in blue (*Alk*) and red (*Eml4*) boxes. Location and sequence of sgRNAs are indicated. PCR primers (gray arrows) show the PCR strategy to identify inversion events.

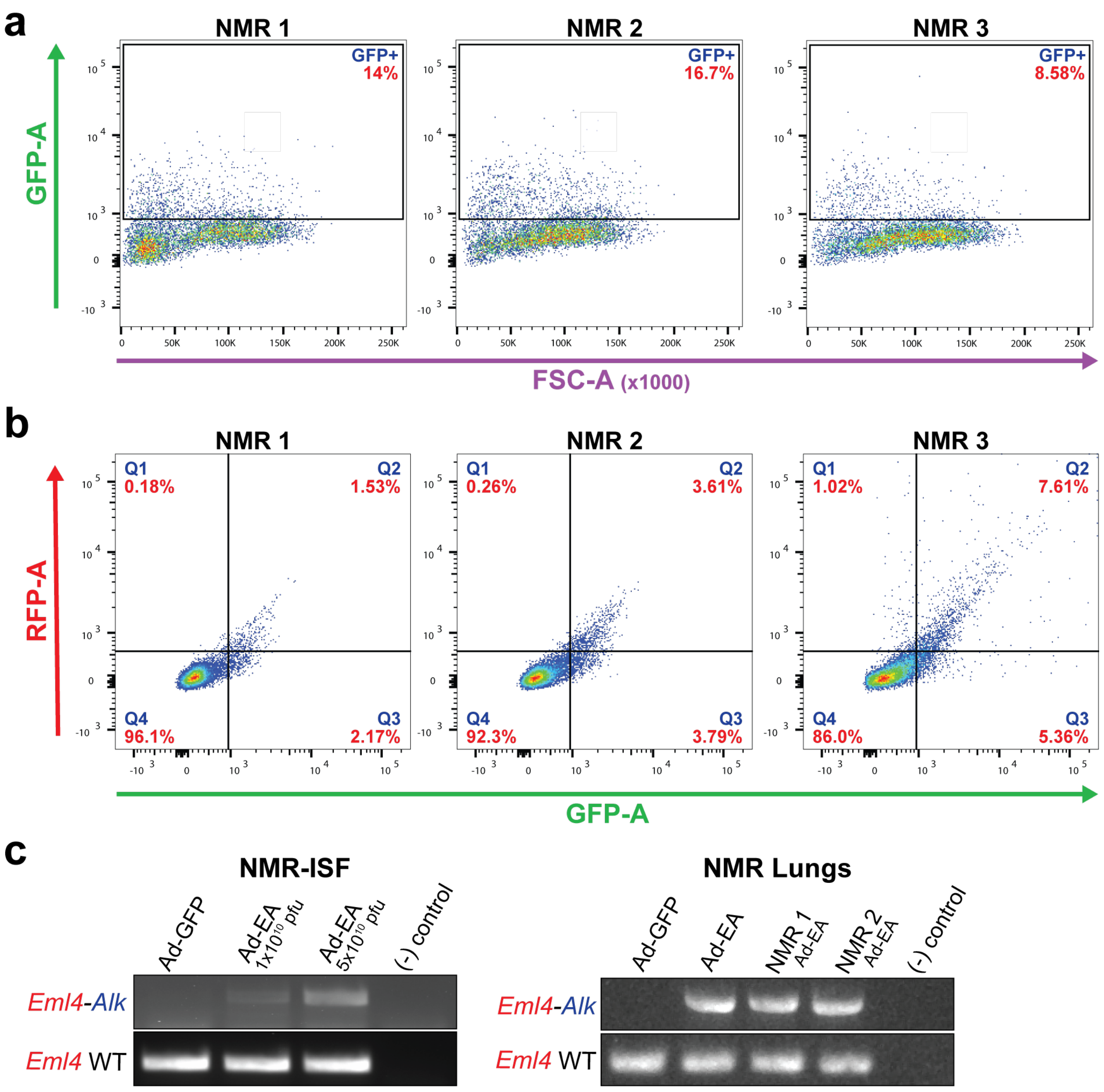


**Fig. S2. Assessing NMR adenovirus infections.** (**a**) NMRs were infected with $3x10^{10}$ pfu of a GFP adenovirus (Ad-GFP; n=3). Lungs were collected after 72 hours and dissociated into single cells for flow cytometry analysis. Percent of GFP-positive cells are indicated in red. (**b**) NMRs were dual-infected with Ad-GFP and Ad-RFP at $3x10^{10}$ pfu total (n=3). Lungs were collected after 72 hours and dissociated into single cells for flow cytometry analysis. Percent of GFP-positive cells are indicated in red, where Q2 indicates dual-positive cells. (**c**) NMR immortalized skin fibroblasts (NMR-ISF) were infected with low ($1x10^{10}$ pfu) and high ($5x10^{10}$ pfu) titer of *Eml4-Alk* adenovirus (Ad-EA). NMR lungs were infected with $3x10^{10}$ pfu of Ad-EA (n=2). Genomic DNA was collected 72 hours after infection. Ad-GFP was used as a negative control. (-) indicates no template control.

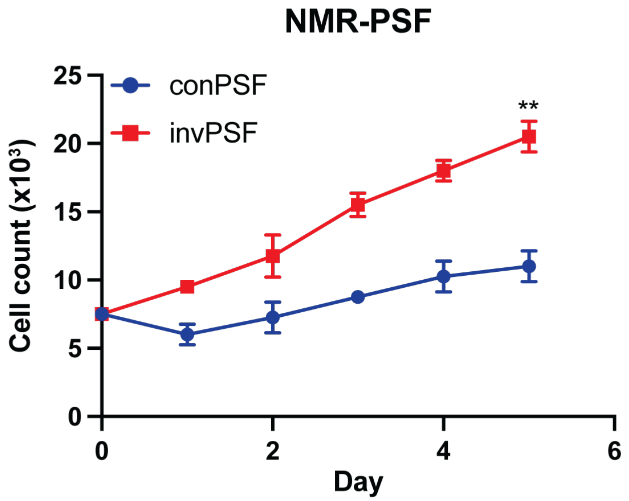


**Fig. S3. Growth assay for control vs. *Eml4-Alk* primary skin fibroblasts.** Cells were plated in triplicate in a 12-well plate, 7,500 cells per well and counted over five days. Significance was calculated using two-way ANOVA and post hoc multiple comparison analysis. Growth assays shown are representative of three independent experiments. Cell lines chosen are representative of one of two cell lines made with the *Eml4-Alk* inversion. Control = conPSF; *Eml4-Alk* = invPSF. Two-way ANOVA determined the interaction of time and genotype to have a statistically significant effect on cell count ($F(5, 20) = 22.70$; p-value $< 0.0001$). Genotype alone also had a statistically significant effect on cell count (p-value = 0.0001). Multiple comparisons determined cell counts on day 5 to be significantly different (**, p-value = 0.0032).

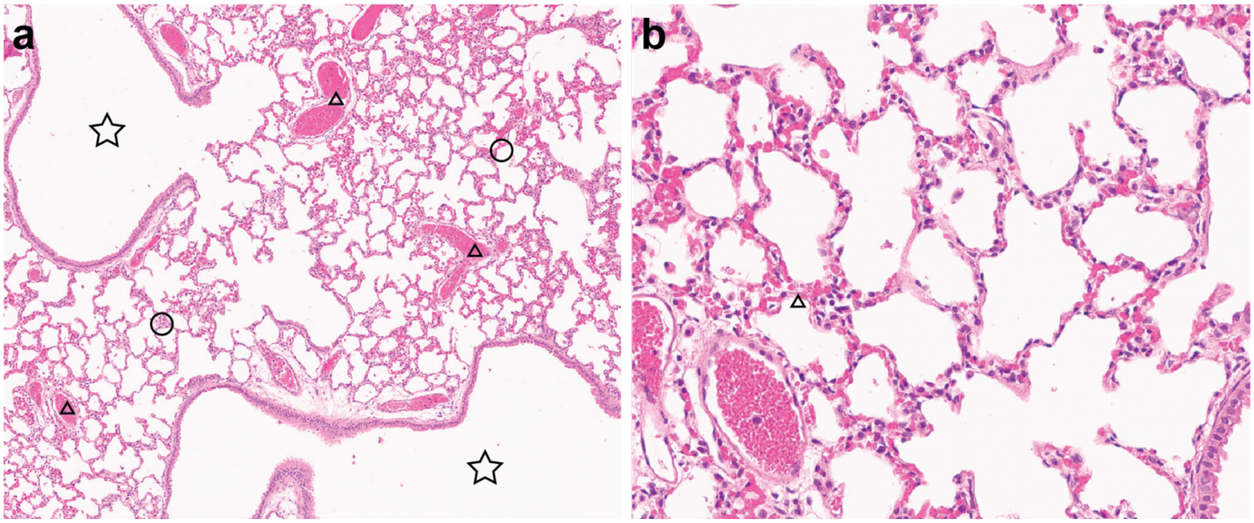


**Fig. S4. H&E analysis of Ad-EA infections at 6-months.** NMR lungs from infections were collected, preserved through FFPE, and processed for H&E staining. Representative images of lungs from the Ad-EA 6-month timepoint (n=5). (**a**) 50X magnification. Star = airway; circle = normal lung tissue; triangle = blood vessel. 50X magnification. (**f**) 200X magnification.

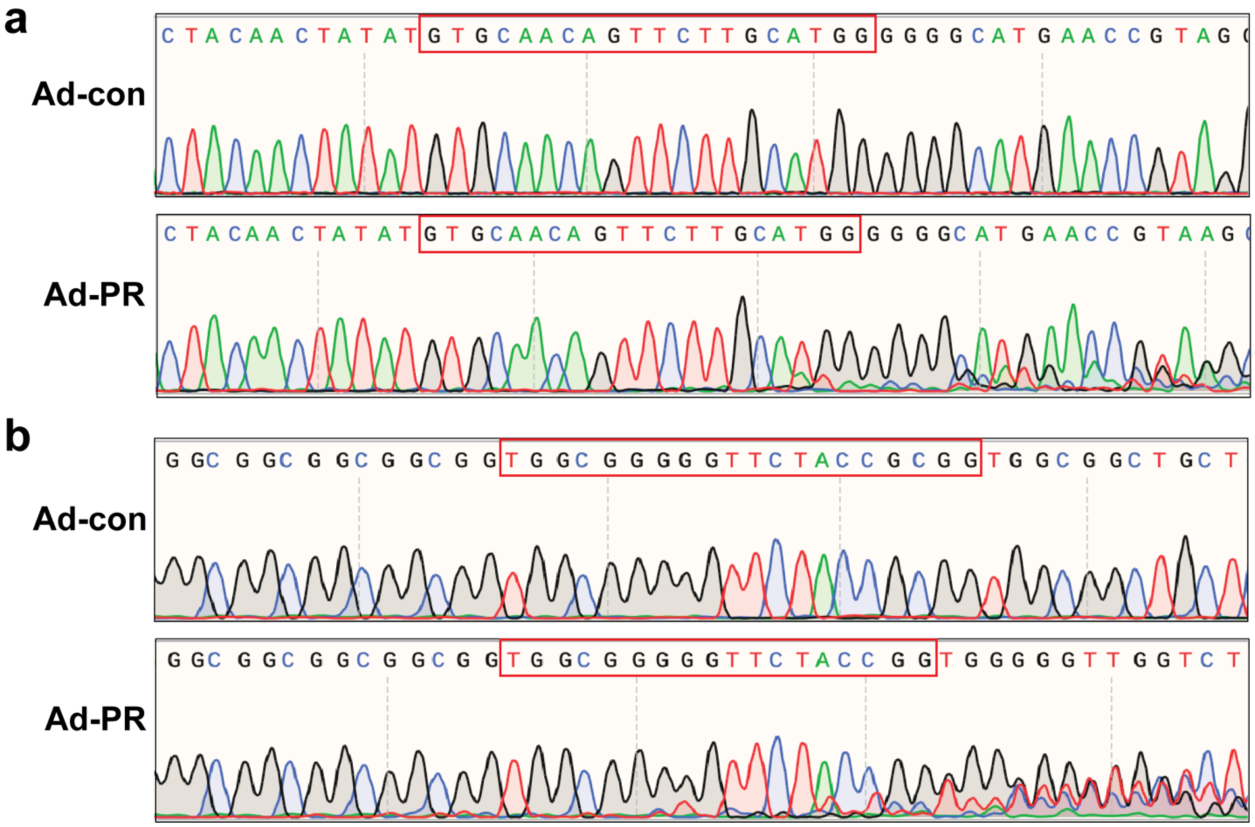


**Fig. S5. Assessing genomic cleavage using the first-generation *Tp53* and *Rb1* adenovirus**. Primary NMR skin fibroblasts were infected with adenovirus containing sgRNAs for *Tp53* and *Rb1* (Ad-PR). Empty adeno-Cas9 (no sgRNAs, Ad-con) was used as a control. DNA was collected, regions surrounding sgRNAs PCR amplified, and sequenced. Red boxes indicate location of sgRNAs. (**a**) Amplification of *Tp53* sgRNA region. (**b**) Amplification of *Rb1* sgRNA region.

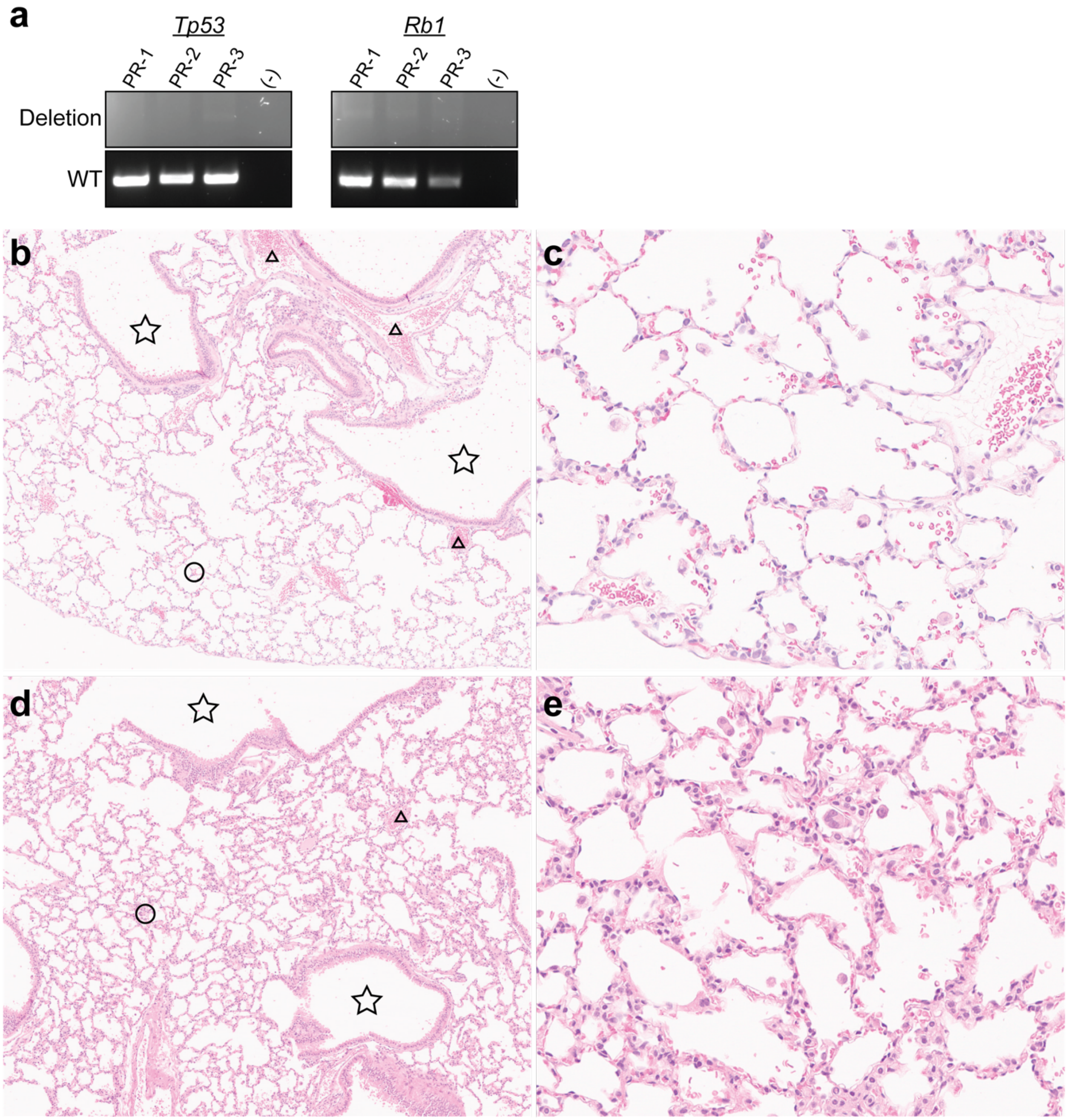


**Fig. S6. PCR and H&E analysis of *in vivo* infections using second-generation *Tp53* and *Rb1* adenoviruses.** (**a**) NMRs were co-infected with Ad-p53-d and Ad-Rb1-d ($3x10^{10}$ pfu). Lungs were collected after 72 hours. DNA was extracted and analyzed for *Tp53* and *Rb1* deletions using genomic PCR. (**b-e**) Infected NMR lungs were collected, preserved through FFPE, and processed for H&E staining. Representative images of lungs from NMR *Eml4-Alk/Tp53* (**b-c**; n=9) and *Eml4-Alk/Rb1* (**d-e**; n=5) infection groups at a 15-week timepoint. Star = airway; circle = normal lung tissue; triangle = blood vessel. (**b**) Representative image for *Eml4-Alk/Tp53* infection, 50X magnification. (**c**) Representative image for *Eml4-Alk/Tp53* infection, 200X magnification. (**d**) Representative image for *Eml4-Alk/Rb1* infection, 50X magnification. (**e**) Representative image for *Eml4-Alk/Rb1* infection, 200X magnification.
